# Repeated cigarette smoke exposure attenuates the proliferation rate of primary human lung fibroblasts *in vitro*

**DOI:** 10.1101/2024.11.27.625673

**Authors:** EC Cannard, A Kronseder, S Karrasch, K Kahnert, D Nowak, M Lehmann, O Holz, MG Stoleriu, RA Jörres

**Affiliations:** Institute and Outpatient Clinic for Occupational, Social and Environmental Medicine, LMU Hospital, Comprehensive Pneumology Center Munich (CPC-M), German Center for Lung Research (DZL), Munich, Germany; Department of Medicine V, LMU Hospital, Comprehensive Pneumology Center Munich (CPC-M), German Center for Lung Research (DZL), Munich, Germany; MediCenterGermering, Germering, Germany; Institute for Lung Research, Lehmann Lab, Biomedical Research Center, Philipps-Universität, Marburg, Germany; Fraunhofer ITEM, Member (BREATH) of the German Center for Lung Research (DZL), Hannover, Germany; Institute for Lung Health and Immunity and Comprehensive Pneumology, CPC-M bioArchive; Helmholtz Zentrum Munich, German Center for Lung Research (DZL), Munich, Germany; Division of Thoracic Surgery Munich, LMU Hospital, Germany; Asklepios Medical Center, Gauting, Germany

**Keywords:** CSE, repeated cigarette smoke exposure, primary human lung fibroblasts

## Abstract

**Background:** Lung fibroblasts are crucial structural cells involved in airway remodelling and tissue maintenance. The study aimed to analyze the effect of cigarette smoke extract (CSE) on primary human lung fibroblasts at various stages of proliferation to assess whether an initial exposure could exert a protective effect against subsequent exposures.

**Methods:** We examined the effect of CSE on fibroblasts isolated from 15 donors who had undergone lung surgery. Fibroblast cultures were exposed to 2% CSE early (day 5-7), late (day 9-11), or at both time points (double). Cell counts and viability tests were performed every 2 days. Changes in proliferation rates were determined immediately at the end of each 2-day exposure as well as 2 days afterwards, until the end of the experiment with a plateau of proliferation. Differences between exposures were analyzed by comparing proliferation rates by ANOVA with adjusted *post-hoc* analyses.

**Results:** Early-, late- and double-CSE exposed cultures showed lower proliferation rates (p<0.05 each) compared to unexposed cultures. The response to late-CSE exposure varied depending on whether an initial CSE exposure had occurred, suggesting a protective effect from the early exposure. Changes observed over the 2-day exposure period and the subsequent 2 days (a total of 4 days) yielded similar results to those recorded during just the 2-day exposure period. Proliferation rates varied among cells from different donors, but responses correlated across different exposure regimens, indicating inherent differences in susceptibility to CSE.

**Conclusions:** Repeated CSE exposures attenuated the proliferation of primary human lung fibroblasts more strongly than a single exposure, despite the fact that an early CSE exposure appeared to protect against the CSE in repetitively exposed fibroblast cultures. Our results highlight that while significant differences may occur among primary cells from different donors in vitro, responses can remain consistent across various exposure conditions.

## Introduction

Chronic obstructive pulmonary disease (COPD) is an irreversible disorder characterized by progressive bronchial obstruction and alveolar destruction causing lung function impairment. Being one of the leading causes of morbidity and mortality worldwide^1,2^, COPD is known to be primarily caused by inhaled noxious agents such as cigarette smoke^3,4^ or particles in ambient air^5^. These agents can provoke senescence of resident cells and induce alveolar destruction and airway remodelling. Based on the link between exposure and biological aging^6–10^, recent studies even highlighted potential beneficial outcomes of drugs with anti-aging effects in ameliorating disease progression in COPD^11,12^.

To mimic hallmarks of COPD such as aging *in vitro*, cell lines and human primary cells have been experimentally exposed to cigarette smoke extract (CSE). The markers of cellular senescence induced by this approach^13–18^ correlated with biochemical and morphological alterations known to occur in lung emphysema^7,8,16,19^.

Lung fibroblasts are structural cells that play crucial roles in airway remodelling and tissue maintenance through their proliferation and regeneration potential^20,21^. This potential is, however, reduced in COPD and in particular in lung emphysema, where alveolar destruction is accompanied by senescence of lung fibroblasts^22,23^. These findings were confirmed in CSE-exposed fibroblast cultures *in vitro*^13,14^. In the majority studies, a single CSE exposure was used to assess cellular responses. This does not address the potential persistence of effects and memory effects of one exposure on subsequent exposures, which might be relevant in view of the fact that in reality smoking occurs as a persistent habit and not a single event.

There are several indicators of cellular senescence and biological aging, including biochemical and molecular markers such as telomere length. The panel of indicators also includes the basic determination of cellular proliferation rate and capacity, which can give direct hints on a potential loss of functionality. Indeed, proliferation rate is closely linked to other markers of senescence and has been demonstrated to be informative particularly for lung emphysema^22–24^.

Based on these considerations, the present study determined, whether the timing of cigarette smoke exposure within the natural course of proliferation of primary human lung fibroblasts resulted in different responses, and whether the response at a later time point would be modulated by a previous exposure, indicating memory effects. To address these questions, we used single or double CSE exposures *in vitro*. In addition, we assessed, to which extent responses quantitatively differed between fibroblasts from different human donors and whether responses would be consistent or inconsistent when comparing exposures.

## Material and Methods

### Patient characteristics

Lung tissue samples were derived from n = 15 patients who had undergone surgery for lung cancer at the Thoracic Surgery Department of Pulmonary Hospital Großhansdorf, Schleswig-Holstein, Germany. All patients gave their written informed consent. The use of the samples for cell culture had been approved by the local Ethics Committee (Ärztekammer Schleswig-Holstein, Bad Segeberg). All patients except two were smokers or ex-smokers. Mean age (range) was 61.6 (45-75) years, 40% were female. Two had a diagnosis of COPD^25^, showing a ratio of forced expiratory volume in 1s to forced vital capacity (FEV_1_/FVC) <0.7 and values of FEV_1_ of 92 and 65 % (predicted); none had severe COPD, as this would have been a contraindication for surgery.

### Isolation of primary human lung fibroblasts

Isolation of fibroblasts from fresh lung samples was performed as previously described^22,23^. Accordingly, peripheral, noncancerous tissue samples derived from the resected lung tissue were stored in Dulbecco’s Modified Eagle Medium (DMEM, gibco, REF:22320-022, Life Technologies Limited) during shipment in a thermal insulation package. After arrival, tissue samples were washed three times in Hanks’ Balanced Salt Solution (HBSS, gibco, REF:14175-053, Life Technologies Limited), pleura and vessels were resected and samples were cut into small pieces which were cultured in DMEM medium supplemented with 10% Fetal Bovine Serum (FBS, gibco, REF:10270-106, Life Technologies Limited), 100 U Penicillin/ml, 100 µg Streptomycin/ml (Penicillin-Streptomycin, gibco, REF:15140-122, Life Technologies Limited) and 0.05 mg/ml Gentamicin (gibco, REF:15710-049, Life Technologies Limited). Fibroblasts were seeded on 24-well plates and incubated at 37°C, 5% CO_2_ and 95% air humidity. To preserve cellular phenotype, only cells at low passage (<4), cultured for about 3 weeks were used for the exposure experiments.

### Preparation of cigarette smoke extract (CSE)

For CSE preparation, a gas washing bottle with a fritted glass (Impinger, DESAGA/Vertrieb Sarstedt AG) filled with DMEM was used, as previously described^26^. Briefly, “Gauloises blond blue” cigarettes with Filter (nominal tar content 10 mg, nicotine 0.8 mg) were smoked during 5 min, while the main stream smoke was conducted through the bottle filled with 10 ml DMEM media. The suction rate was set so that each cigarette was smoked during 5 min to a length of about 5 mm from the cigarette filter. The generated CSE was sterile filtered, aliquoted and stored at -32°C. Freshly thawed CSE aliquots (100% reference) were used for all experiments and diluted to the final exposure concentration of 2%.

### Proliferation assay

Before reaching full confluence, cells were transferred to 24-well plates for the CSE exposure. To avoid potential topographical biases on the well plates, one inside well and one outside well were identically treated and the average of both cell counts was taken for analysis. For each time point a separate plate was prepared.

To analyze cell proliferation differences over time, 4 groups were designed: control (ctrl, only medium without CSE), single early CSE exposure (se-CSE), single late CSE exposure (sl-CSE) and early plus late CSE exposure (double-CSE). Cells were counted every second day starting with the day 3 post-seeding. Exposure to se-CSE was performed between day 5 and 7, sl-CSE exposure between day 9 and 11, double-CSE at both time points. In total, cells were counted 9 times over a time period of 18-19 days. Proliferation rates were determined at the end of the exposure (2-days period) and 2 days thereafter separately. These results were also combined into a 4-day rate. The experimental approach is summarized in **Figure 1A**.

**Figure 1.**
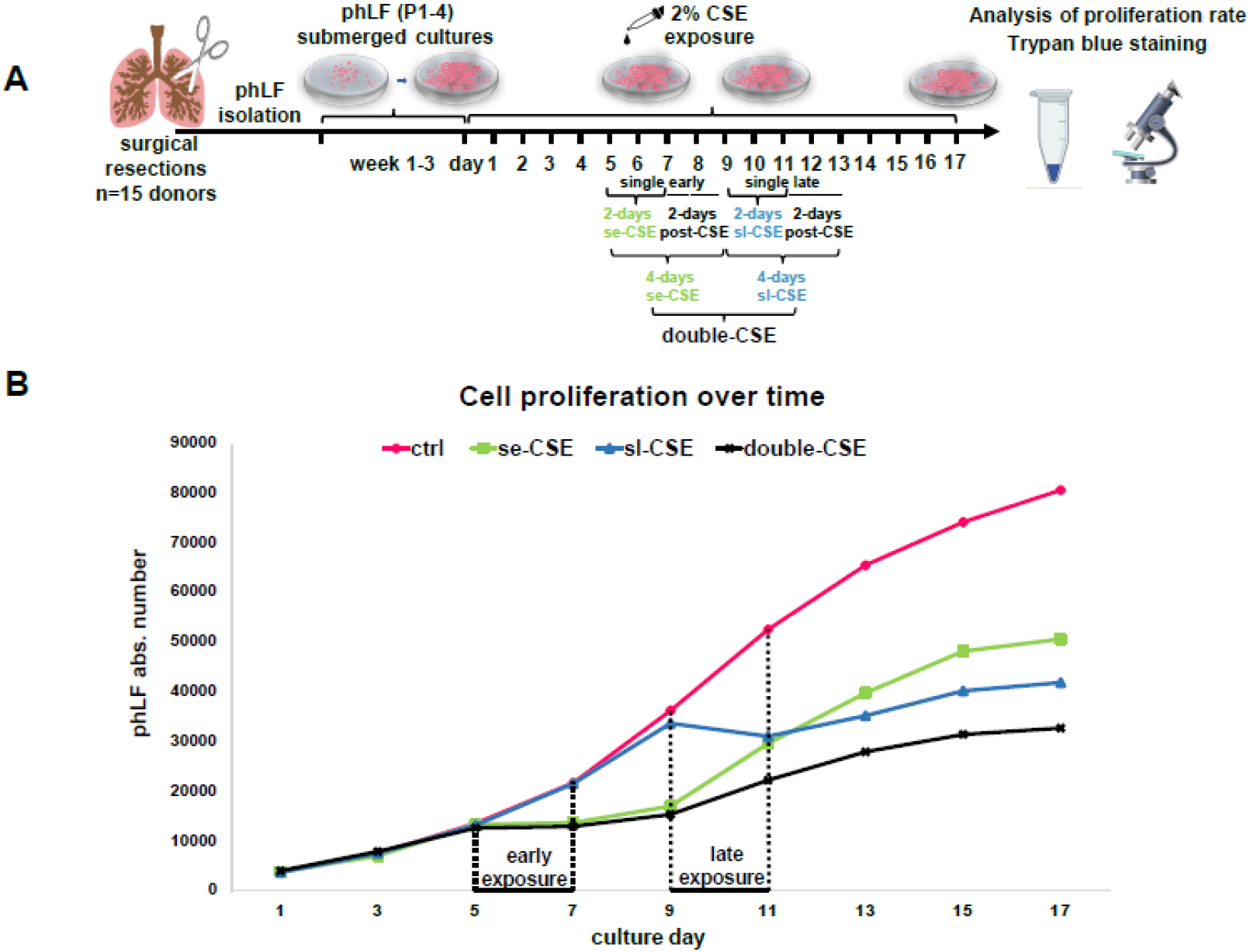
**Panel A:** Overview of the experimental approach. **Panel B:** Mean values of proliferation curves (see **Table 1**) based on the absolute cell number of primary human lung fibroblasts at different time points and upon single early, single late or double-CSE exposure. For statistical results see text. P1-4: passage 1-4, phLF: primary human lung fibroblasts, CSE: cigarette smoke extract, ctrl: control, se-CSE: single early CSE exposure, sl-CSE: single late CSE exposure

### Cell count

Cells were harvested using trypsin and counted in a Neubauer-chamber by light microscopy, according to the previously described protocol^22^. Viability was evaluated using trypan blue staining. The average of three counts was taken.

**Table 1.**
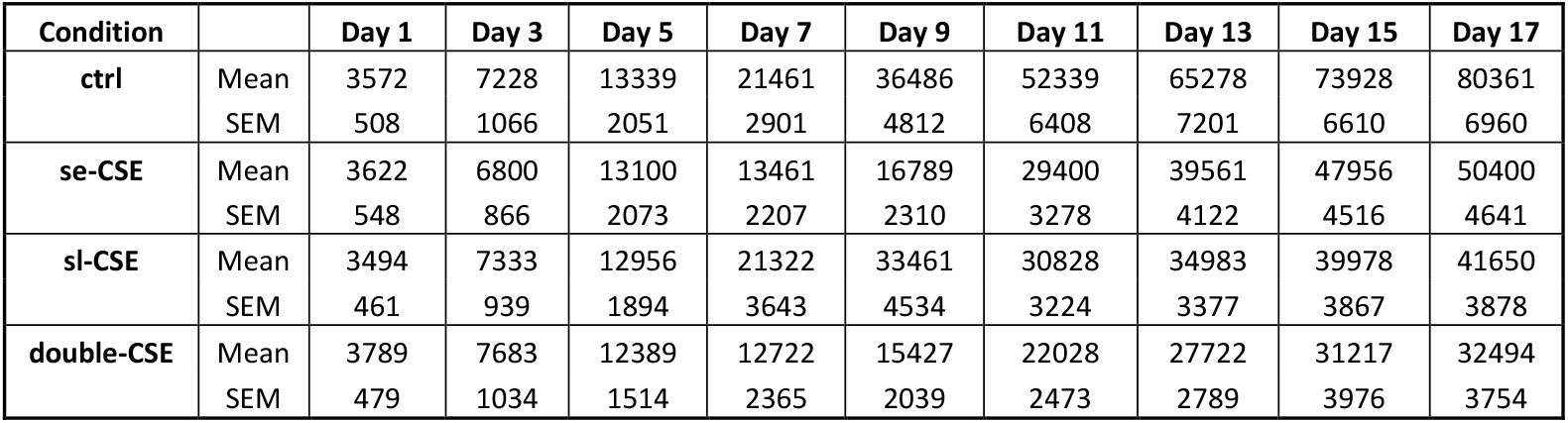
Absolute numbers of fibroblasts at different time points of culture. Values are given as mean and SEM (standard error of mean). ctrl: control exposure (medium), se-CSE: single early CSE, sl-CSE: single late CSE, double-CSE: early and late CSE exposure, CSE: cigarette smoke extract

### Repeated exposure study

Exposures were performed with 2% CSE. In the control group, cells received the same amount of culture medium without CSE. Each experiment used cells from a different donor in duplicate. Each exposure lasted 2 days, as in the single exposures, and was stopped by washing the cells 3 times with DMEM. For an overview see **Figure 1A**.

### Data analysis

Changes in proliferation rate were calculated between defined time points, either immediately after the 2-day exposure versus immediately before exposure, or 2 days after the end of exposure versus immediately before exposure (therefore comprising 4 days), or immediately after exposure versus 2 days after the end of exposure. The primary changes used in the present analysis were those covering the 4 days from the start of exposure until 2 days after the end of exposure. The cell numbers determined at these time points were used to compute the respective ratios of cell numbers as numerical values of the respective proliferation rates. In addition, ratios between proliferation rates for different exposure conditions were calculated, if adequate. All ratios were log_10_-transformed for analysis in order to achieve normal distribution. The results were re-transformed to obtain geometric mean values with geometric standard errors of mean (SEM), whereby geometric SEM is to be understood as variability factor of the geometric mean. Moreover, 95% confidence intervals were computed for the geometric mean values. Statistical comparisons between exposures were performed by the Wilcoxon matched-pairs signed-ranks test, and p-values are given explicitly; thus, no correction for the multiplicity of tests was introduced. For correlation analysis, Pearson’s correlation coefficients of log-transformed data were computed. p-values < 0.05 were considered as statistically significant. All statistical analyses were performed using the software package SPSS (Version 26., IBM, Armonk, NJ, USA).

## Results

### Cellular proliferation curves over time

CSE exposure reduced the slope of the fibroblast proliferation curves after se-CSE, sl-CSE and double-CSE compared to ctrl (**Figure 1B**). Numerical values can be found in **Table 1**.

The number of trypan blue-negative cells was proportional to the total cell number, while the number of trypan blue-positive cells was very low at all time points and showed no significant differences between exposure conditions.

### Relative changes in cell number over 4 days

#### Early exposure

To quantify effects of early CSE exposure, data of the se-CSE and the first of the double-CSE exposures could be combined as mean values. Similarly, as a reference for the early time point, proliferation rates of ctrl cultures and sl-CSE could be combined as mean value. When using these combined values, the post/pre-CSE ratio (day 9/5) of proliferation rates was significantly lower with CSE in comparison to ctrl (geometric mean (geometric SEM) 1.25 (1.06) versus 2.72 (1.08), p = 0.001, **Figure 2A**). Correspondingly, early CSE exposure reduced the proliferation rate compared to ctrl to about half its value (geometric mean (geometric SEM) 0.46 (1.06), 95% CI: 0.40; 0.52).

**Figure 2.**
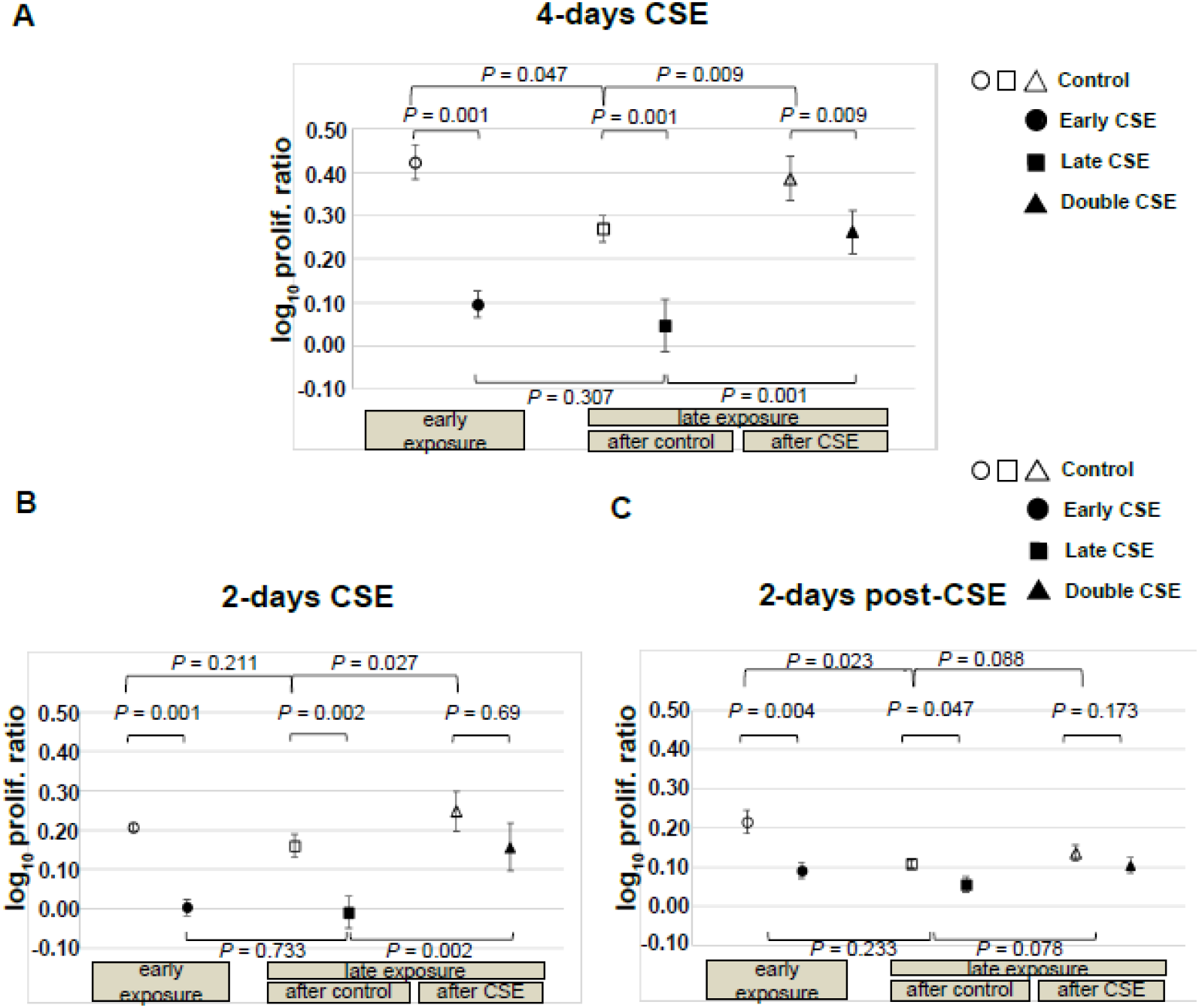
**Panel A:** Relative changes in cell number (expressed as log_10_) of fibroblasts over the 4-day period between the start of exposure and 2 days after exposure, distinguishing between different early and late CSE exposures. For further understanding, the changes over 4 days were split into the changes over consecutive 2-day periods as shown in the lower panels. **Panel B:** Relative changes in cell number during the 2-days period between the start and the end of exposure. **Panel C:** Relative changes between the end of exposure and 2 days after exposure. Comparisons between conditions were performed by the Wilcoxon matched-pairs signed-ranks test. Please note that all values are depicted as log_10_ of ratios of proliferation rates (prolif. ratio), which can be reconverted into ratios by taking them as power to base 10. These reconverted values equivalent to geometric mean values and the corresponding geometric standard errors of mean are given in the text.

#### Late exposure

In the subsequent analyses, we determined the effect of different early exposures on the responses observed in late exposures to reveal potential protective or enhancing effects of early CSE exposure.

##### Comparisons involving similar exposure histories

If early exposures did not involve CSE, the effect of a single late CSE exposure could be determined. This also showed a significant effect, as ctrl cultures showed higher relative proliferation rates (1.86 (1.08)) compared to sl-CSE exposed cultures (1.11 (1.14), p = 0.001, **Figure 2A**). Correspondingly, in terms of geometric mean values the post/pre-CSE ratio (day 13/9) in comparison to ctrl was 0.60 (1.10) with 95% CI: 0.49; 0.73.

If early exposures involved CSE, the effect of the late of the double CSE exposure could be determined. This again revealed a significant effect, as se-CSE cultures showed higher relative proliferation rates (2.43 (1.12)) compared to double-CSE exposed cultures (1.83 (1.13), p = 0.009, **Figure 2A**). Accordingly, in terms of geometric mean values the post/pre-CSE ratio (day 13/9) in comparison to se-CSE was 0.75 (1.08) with 95% CI: 0.63; 0.89.

##### Comparisons involving different exposure histories

To determine the effect of early CSE exposure on the late CSE exposure, the late of the double-CSE exposure was compared with the sl-CSE exposure that was not preceded by early CSE exposure. The late of the double exposure showed a significantly lower reduction of proliferation rate compared to the sl-CSE (p = 0.001, **Figure 2A**). Therefore, the effect of CSE in the late exposure was lower when CSE had already been administered in the early exposure, suggesting a protective effect.

In a similar manner, it could be determined whether the proliferation rate at the time of the second exposure but without CSE, depended on early exposure to CSE. In this case, the rate was significantly reduced if CSE had been given previously (p = 0.009), indicating a persistent effect of the early CSE exposure.

##### Separate analysis of effects observed during and after exposures

Regarding the responses during the 2 days of exposure, very similar findings as for the 4-day periods were observed (**Figure 2B**). The results obtained for the 2-days after exposures were also similar, although proliferation rates were much lower and differences between exposure conditions smaller (**Figure 2C**).

#### Correlation analyses

#### Early exposure

For this purpose, we analyzed the individual 4-day exposure differences for each donor (**Figure 3**). In the first step, it was assessed whether baseline proliferation rates were reproducible. Indeed, when comparing ctrl and sl-CSE exposed cultures at the early time point (day 5-9), proliferation rates showed a significant correlation between the two conditions (r = 0.724, p = 0.002; **Figure 3A**), with data scattering around the line of identity. Moreover, mean values for the two conditions were not significantly different (Wilcoxon test, p = 0.733). This indicated that individual baseline proliferation rates were reproducible.

**Figure 3.**
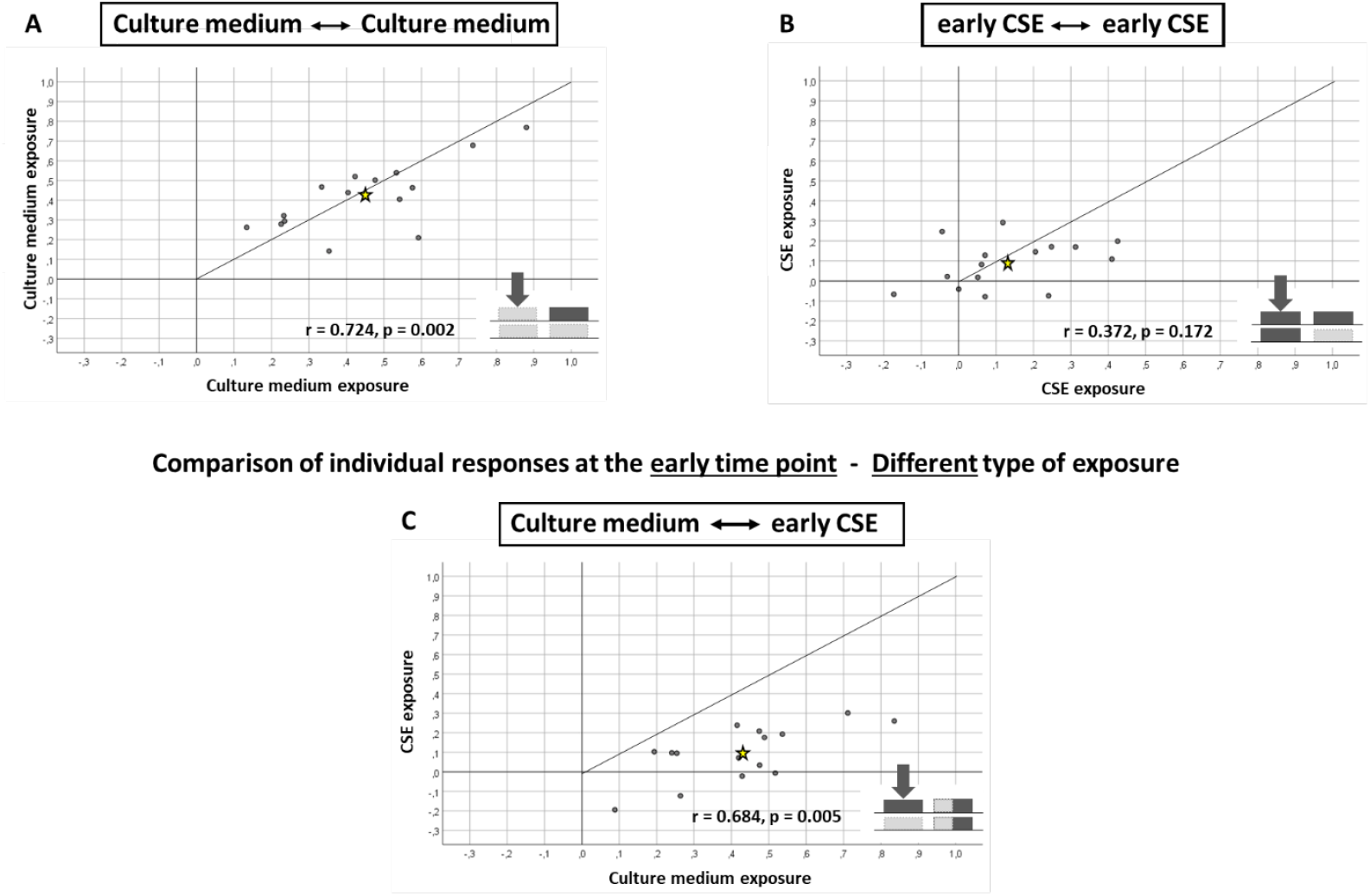
Correlation analysis between the changes in proliferation rates of primary lung fibroblasts derived from 15 donors at the early time point. For quantification, Pearson’s correlation coefficient (r) was computed. The star indicates the mean values. Additionally, lines of identity are shown. Data are shown as log_10_ of the ratio of proliferation rates (prolif. ratio) over 4 days (2 days after exposures compared to values before the 2-day exposures). **Panel A:** Response to culture medium in the first exposures of either the ctrl or the sl-CSE condition. **Panel B:** Response to CSE in the first exposures of either the se-CSE or the double-CSE condition. **Panel C:** Response to CSE versus culture medium in the first exposures, using mean values of the se-CSE and double-CSE conditions versus mean values of the ctrl and sl-CSE conditions.

To determine to which extent the responses to the early CSE exposure were reproducible, the respective data from the se-CSE and double-CSE exposures were used. In this case, individual proliferation rates did not significantly correlate with each other (r = 0.372, p = 0.172; **Figure 3B**). There were also no significant differences between the two conditions (p = 0.307). This indicated that reproducibility of early CSE responses was less than that of baseline proliferation.

On the other hand, there was a positive correlation between baseline proliferation and proliferation after CSE exposure for the early time point (r = 0.684, p = 0.005; **Figure 3C**), again using mean values of the corresponding exposures (see above). Of course, mean values showed a significant difference between the two conditions (p = 0.001). Taken together, these observations suggest that the intraindividual differences of the responses between CSE and control exposure are better reproducible than the absolute values of the responses, again underlining intrinsic differences between cells of different donors.

#### Late exposure

In the next step, reproducibility of late responses was analyzed if, previous history of early exposures war either the same or different (**Figure 4**).

**Figure 4.**
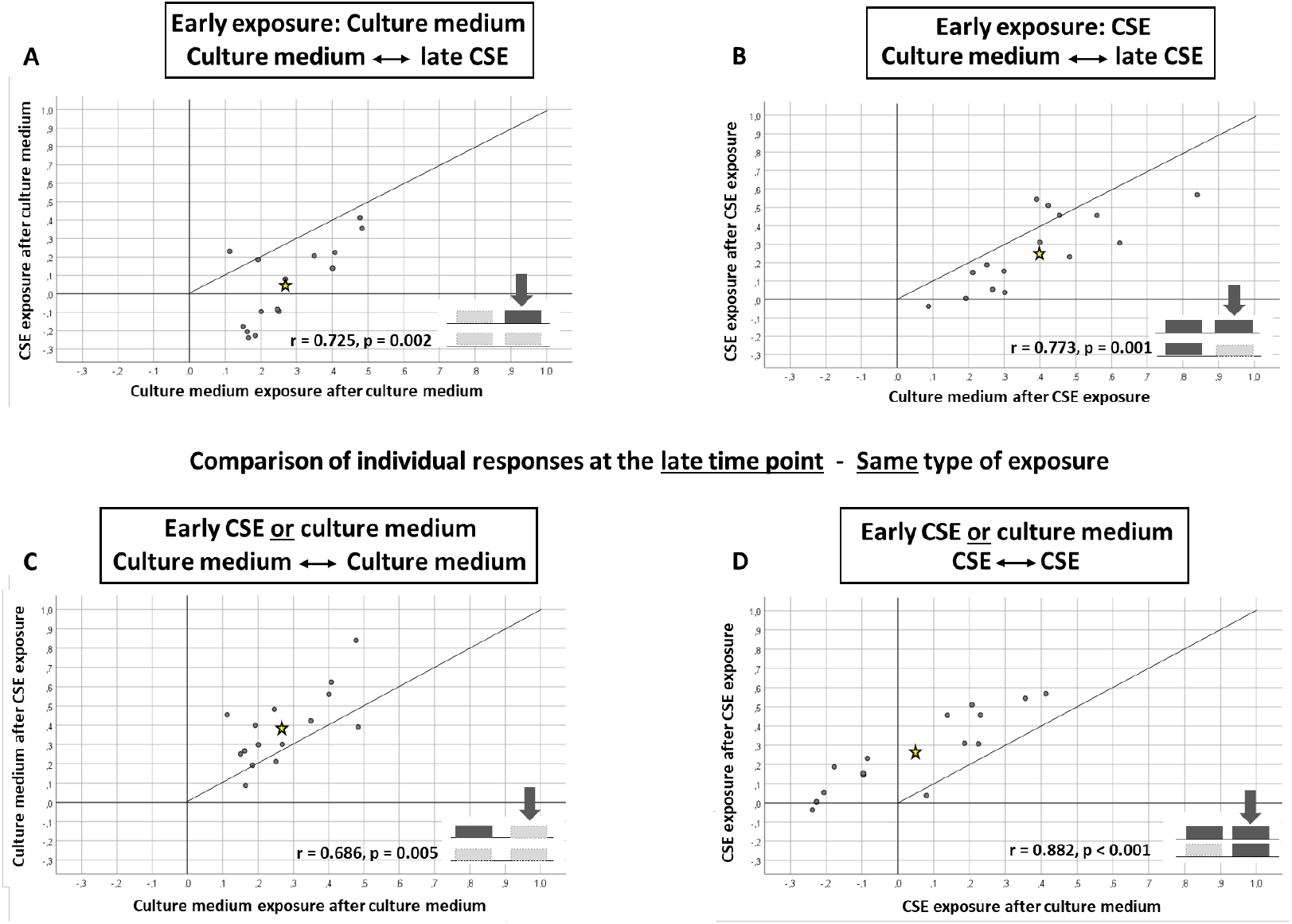
Correlation analysis between the changes in proliferation rates of primary lung fibroblasts derived from 15 donors at the late time point. For quantification, Pearson’s correlation coefficient (r) was computed. The star indicates the mean values. Additionally, lines of identity are shown. Data are shown as log_10_ of the ratio of proliferation rates (prolif. ratio) over 4 days (2 days after exposures compared to values before the 2-day exposures). **Panel A:** Late CSE vs. culture medium exposure after early exposure to culture medium. **Panel B:** Late CSE vs. culture medium exposure after early exposure to CSE. **Panel C:** Late culture medium exposure when early exposures involved either culture medium or CSE. **Panel D:** Late CSE exposure when early exposures involved either culture medium or CSE. From the correlation coefficients it can be seen that the responses were fairly reproducible across the fibroblasts from different donors.

When comparing the sl-CSE with the ctrl condition, proliferation rates for the second exposure without previous CSE exposure could be analyzed. These rates correlated with each other (r = 0.725, p = 0.002; **Figure 4A**), again demonstrating lower proliferation for CSE compared to culture medium (p = 0.001).

When exposure history comprised an early CSE exposure, the response to CSE at the late time point also correlated with the response to medium (double-CSE vs. se-CSE condition, r = 0.773, p = 0.001; **Figure 4B**). The significant difference between mean values (p = 0.009) indicating lower proliferation for CSE exposure at the late time point.

To complete the correlation analysis, we also compared responses when cultures had different early exposure histories (culture medium or CSE). When late exposures included the administration of culture medium, responses correlated significantly with each other (se-CSE vs. ctrl condition, r = 0.686, p = 0.005; **Figure 4C**), and mean values were significantly different (p = 0.009). When the second exposure involved CSE, the results also showed a positive correlation (double-CSE vs. sl-CSE condition, r = 0.882, p < 0.001; **Figure 4D**), and proliferation rates were also significantly different (p < 0.001), again illustrating the protective effect of a previous CSE exposure.

## Discussion

Our study revealed that twofold CSE exposure attenuated proliferation rate in primary human lung fibroblasts stronger than a single exposure. Accordingly, CSE exposure induced a flattening of proliferation curves in cell culture, in line with previous results obtained with immortalized fibroblast cell lines^13^. On the other hand, early CSE exposure reduced the relative effect of CSE in a second exposure, indicating memory or even protective effects of the first exposure. Our data also demonstrated remarkable differences in the proliferation rates of fibroblasts derived from different donors. Irrespective of this, the relationship between the responses to either CSE or culture medium was reproducible for most of the exposure conditions. This suggests that the observed effects including the attenuation of a response to CSE by a previous CSE exposure were generic effects in human lung fibroblasts and not specific for one cell line. The results also suggest that repeated exposures in cell culture might lead to different outcomes compared to single exposures, although in principle inhibitory effects of CSE could be elicited over different time points in the natural course of cell proliferation. This observation may be relevant for drawing conclusions from CSE experiments on the pathophysiological mechanisms *in vivo*, as COPD patients present a long-term smoking history that may not be adequately modelled by single CSE exposures.

The largest absolute decline in proliferation rates upon single exposure was observed with single late CSE exposure (see **Figure 2**) but the reduction was also strong for single early CSE exposure. Although proliferation rate increased after approximatively 3-5 days after single early exposure, it did not reach the rate and the capacity of the unexposed cells. This behavior was already observed by Kanaji and coworkers and might be explained by the co-existence of fibroblast sub-populations not affected by CSE exposure and capable of partially resuming proliferation^27^.

The strongest reduction in cell number observed with twofold CSE exposure aligns with data from Nyunoya and colleagues^13,14^. They found that a single exposure temporarily inhibited cell proliferation, but a repetitive exposure irreversibly induced cell arrest and senescence. In addition, previous reports demonstrated persistent effects of even single short-time CSE exposures on primary lung fibroblasts, depending on CSE concentration and confirmed by beta-galactosidase as senescence marker^26,28^. To explain these responses, it has been suggested that repetitive CSE exposures induce two fibroblast phenotypes—senescent and profibrotic—which contribute to irreversible morphological changes in lung emphysema^27^. The authors suggested that upon CSE exposure resilient lung fibroblasts acquire a profibrotic phenotype promoting peri-bronchial fibrosis in COPD^27^.

These considerations are in line with our results showing very low proliferation rates after late compared to early CSE exposures, if one assumes that the induction of these two phenotypes occurs only in cells that are relatively young and far from the final proliferation plateau. A dependence of CSE effects on the time point of exposures might also underly our observation of a relatively lower effect of CSE, if CSE had been administered previously, indicating partial protection against subsequent CSE exposures. The underlying mechanisms were not analyzed in our study but might be related to upregulated anti-oxidant defenses, as oxidants have been shown to play a role in the response to CSE^29^.

The observed response patterns were similar whether evaluated over 2-day exposure periods or 4-day periods that included a 2-day post-exposure period. Proliferation rates were lowest immediately after the 2-day exposures, suggesting that the dominant effect within the 4-day period was due to the acute responses. However, the results for both types of data evaluation were very similar; thus, our choice of the 4-day period for primary data analysis did not affect the conclusions.

A key aspect of our study was the inclusion of primary lung fibroblasts from various donors. This allowed the assessment of differences in the response of different cell lines and to test whether the responses of cell lines were consistent across the four different exposure conditions. This appears of interest as primary cells may differ in their response from established cell lines. On the other hand, the availability of primary cells may be limited and therefore experiments are performed with one or only a few cell lines. That cells from different donors show different degrees of response, is conceivable. It would be disastrous, however, if the relationship between responses to various types of exposures would be different, as this might easily lead to inadequate conclusions.

Indeed, fibroblasts derived from different donors showed different proliferation rates, reflecting the heterogeneity of fibroblast populations and differences in patients’ history, as previously described^30,31^. Importantly, however, the relationship between the responses to different exposure conditions was consistent in nearly all comparisons (see **Figures 3 and 4**), and baseline proliferation rates were reproducible. These observations suggest that even from a small number of primary cell lines reliable conclusions may be drawn.

The present study has several limitations. Since environmental toxin exposure is a long-term process, multiple exposures would be most adequate but may interfere with the limited proliferation capacity *in vitro*. This might be solved by continuous exposures under steady-state conditions. Second, each exposure was stopped by washing the cells in repeated washing steps. Although unlikely, a small amount of CSE might have remained on the cells. Based on previous findings^22,23^, we selected proliferation rate as outcome parameter. This is sensitive and meaningful marker of cellular senescence but of course of limited value for the analysis of mechanisms. At the time of the experiments performed in 2005/6, more advanced methods were not available in the laboratory, due to technical and logistic reasons. Irrespective of this, we believe that the analysis of proliferation rate which is linked to both, cellular senescence and regeneration potential, was already sufficient to provide valuable insights into the effects of cigarette smoke on human fibroblasts and that the results are still of interest.

In conclusion, the present study demonstrated that cigarette smoke extract irreversibly affected the proliferation rate of primary human lung fibroblasts *in vitro* at early and late phases of cell culture. The results suggest a higher susceptibility to cigarette smoke in the late phase but also a relative protective effect of an early exposure on the response to a late-phase cigarette smoke exposure, suggesting the induction of protective mechanisms. Absolute proliferation rates showed marked differences between cells from different donors. Despite this, the relationship of responses to different exposure conditions remained similar across cells, suggesting that the observed effects of cigarette smoke were generic for human lung fibroblasts. Our observations might be of interest for the understanding of COPD and the design of *in vitro* models of this disease.

## Authors’ contributions

RAJ conceptualized the study, ECC, AK performed the experiments, KK, SK collected the clinical data, MGS and RAJ analyzed the clinical data and documentation, ECC and RAJ performed the statistical analysis, RAJ, MGS, DN, KK, OH and ML aided in interpreting results. ECC, MGS, and RAJ drafted and wrote the manuscript. ECC, MGS and RAJ also revised the manuscript. All authors discussed the results and commented on the manuscript. This work was part of the medical doctoral thesis of ECC.

